# Ensemble Post-hoc Explainable AI for Multilead ECG: Identifying Disease-Relevant Features in Single-Lead Interpretations

**DOI:** 10.1101/2025.02.14.638219

**Authors:** Jacqueline Michelle Metsch, Philip Hempel, Miriam Cindy Maurer, Kristin Elisabeth Steinhaus, Nicolai Spicher, Anne-Christin Hauschild

## Abstract

**Background:** Despite the growing success of deep learning (DL) in multivariate time-series classification, such as 12-lead electrocardiography (ECG), widespread integration into clinical practice has yet to be achieved. The limited transparency of DL hinders clinical adoption, where understanding model decisions is crucial for trust and compliance with regulations such as the General Data Protection Regulation (GDPR) or the EU AI Act.

**Results:** To tackle this challenge, we implemented a widely used 1D-ResNet in Pytorch that was trained on the large-scale Brazilian CODE dataset to classify six different ECG abnormalities. We employed the model on the German PTB XL dataset, and evaluated its decision-making processes using 16 post-hoc explainable AI (XAI) methods. To assess the clinical relevance of the model’s attributions, we conducted a Wilcoxon signed-rank test to identify features with significantly higher relevance for each XAI method. We used an ensemble majority vote approach to validate whether the model has learned clinically meaningful features for each abnormality. Additionally, a Mann–Whitney U test was employed to detect significant differences in relevance attributions between correctly and incorrectly classified ECGs. Overall, the model achieved sensitivity scores above 0.9 for most abnormalities in the PTB XL dataset. However, our XAI analysis showed that the model struggled to capture clinically relevant features for some diseases. Certain XAI methods, including DeepLift, DeepLiftShap, and Occlusion, consistently highlighted clinically meaningful features across abnormalities, while others, such as LIME, KernelShap, and LRP, failed to do so. Moreover, some XAI methods demonstrated significant differences in attributions between correctly and incorrectly classified ECGs, highlighting their potential for enhancing model robustness and interpretability.

**Conclusion:** Our findings underscore the importance of selecting suitable XAI methods tailored to specific model architectures and data types to ensure transparency and reliability. By identifying effective XAI techniques, this study contributes to closing the gap between DL advancements and their clinical implementation, paving the way for more trustworthy AI-driven healthcare solutions.

## 1 Background

Deep learning (DL) models have demonstrated remarkable performance across various domains, including healthcare [28], finance [27], and autonomous systems [53]. However, due to their inherent complexity they are considered “black-box” models, offering little to no transparency regarding their decision-making processes. This lack of explainability is particularly concerning in high-risk applications such as medical diagnosis, where understanding a model’s reasoning is critical for gaining the users’ trust. In particular for medical data, there are multiple examples where DL models learned shortcuts in the data, leading to classifications unrelated to diagnostic features and resulting in misdiagnosis [10]. The recently proposed EU AI Act [1] explicitly underscores the need for transparency and accountability in AI development and application, enforcing the importance of interpretability.

To address these concerns, researchers have increasingly turned to Explainable AI (XAI) methods to analyse whether a model learned the correct features from input data [7, 26, 30]. These techniques aim to provide insights into the internal workings of DL models, helping to validate their predictions and detect biases or spurious correlations. Their effectiveness has already been demonstrated in image classification tasks, where explanations are typically visualized as heatmaps that indicate whether the model focuses on the correct regions of the image [13, 44]. A range of XAI methods, including gradient-based approaches, perturbation-based methods, and model-agnostic techniques, can be employed to assess the decision-making process of deep networks [17]. However, the choice of the XAI method significantly impacts the interpretations derived from a model. [4, 15, 22, 25] found that different XAI techniques can produce varying relevance attributions for the same model and dataset, leading to potential inconsistencies in explainability results. Metsch et al. [25] for example found that IntegratedGradients, DeepLift, DeepShap and GradientShap perform similarly well on different datatypes, while Deconvolution, Guided Backpropagation, and LRP-*α*1-*β*0 differ from other methods. This variability gives rise to concerns about the reliability and biases of XAI-based model validation, underscoring the necessity of selecting appropriate interpretability techniques tailored to the specific application. It is, therefore, crucial to identify these discrepancies and address them accordingly in order to ensure that explainability efforts genuinely enhance trust and model accountability rather than introducing further ambiguity. In feature selection, the issue of method-specific bias has been tackled using ensemble approaches that compute a quantitative, consensus-based relevance score [6, 31]. A similar strategy has emerged in XAI, where ensemble techniques typically aggregate heatmaps produced by multiple explanation methods to mitigate individual biases [12, 57]. Additional feature selection approaches, such as multi-step feature selection [50], were not explored in this study due to time constraints.

In this study, we address the issue of transparency and reliability by developing a comprehensive processing pipeline that integrates three steps for evaluating the applicability of XAI Methods for time series in medicine and proposes a novel ensemble approach. We implement the following steps: (1) a widely used 1D-ResNet was trained on the large-scale CODE dataset, (2) we evaluated its prediction performance on the German PTB XL dataset, (3) benchmarked the decision-making processes using 16 post-hoc explainable AI (XAI) methods, and (4) used an ensemble majority vote approach to validate whether the model has learned clinically meaningful features. Figure 1 summarizes the pipeline.

**Figure 1:**
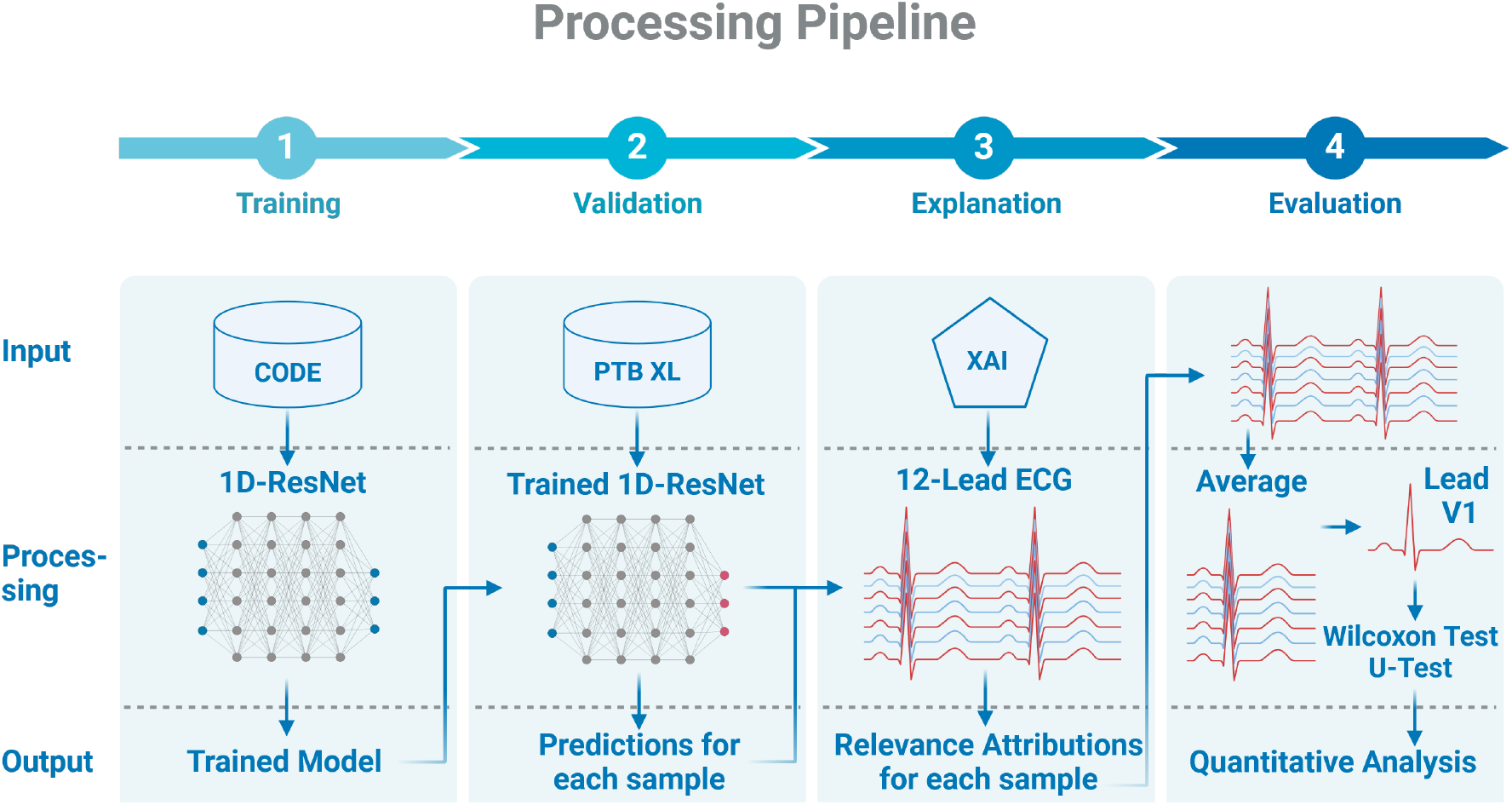
Processing pipeline employed in this paper. First, we train a widely used 1D-ResNet on a large-scale ECG dataset from Brasil. Afterward, the trained model is used to generate predictions for data stemming from the German PTB XL dataset. Subsequently, we apply 16 different post-hoc XAI methods to generate relevance attributions for the predicted samples. Lastly, we developed a processing pipeline that averages the ECGs as well as their relevance attributions, then focuses on the important lead V1 and performs two types of statistical tests for quantitative analysis of the results.

### 2 Methods

ECG is a measurement of the electrical activity of the human heart and standard of care for assessment of cardiovascular health. Typically, 10 cables are connected to the human body, resulting in 12 signals recorded in parallel – entitled “leads” – making ECG signals multivariate time series.

We first train a widely used 1D-ResNet model [34] on the large-scale Brazilian CODE dataset [34–36] to classify six different abnormalities in 12-lead ECG data and then generating explanations using 16 different XAI methods based on the models predictive performance on the German PTB XL dataset [49]. We use a majority vote ensemble approach on the XAI methods to more robustly identify whether the model learned clinically relevant features to predict the different abnormalities. In the realm of feature selection methods, ensemble approaches are already being employed to reduce biases of specific methods [6, 31]. In the field of XAI, ensemble approaches are also gaining popularity, though they often focus on aggregating heatmaps generated by multiple XAI methods [12, 57]. Additionally, we employ a quantitative analysis of all XAI methods to identify which methods are appropriate for our specific classification task, as well as their capabilities in discerning between true positive and false positive classified samples. See Figure 1 for details.

### 2.1 CODE Data

The Brazilian CODE dataset [34–36] comprises multiple 12-lead ECGs from 1, 676, 384 unique patients (mean age: 51.2; 60.3% female), recorded through the Telehealth Network of Minas Gerais (TNMG). Telehealth refers to a remote healthcare model in which specialists (e.g., cardiologists) virtually review ECGs sent from primary care facilities, ensuring high-quality diagnoses despite geographical distance. The ECGs were acquired between 2010 and 2016 in 811 counties, with each 7–10-second trace is annotated for six key abnormalities: (1) 1st degree AV block (1dAVb), (2) Complete Right Bundle Branch Block (CRBBB), (3) Complete Left Bundle Branch Block (CLBBB), (4) Sinus Bradycardia (SB), (5) Atrial Fibrillation (AF), and (6) Sinus Tachycardia (ST). These specialist-reviewed labels provide a robust ground truth while making CODE one of the largest and most comprehensive labeled 12-lead ECG repositories available for developing and evaluating AI models. The CODE Study was approved by the Research Ethics Committee of the Universidade Federal de Minas Gerais (protocol 49368496317.7.0000.5149)

### 2.2 PTB XL Data

PTB XL [49] contains 21799 clinical 12 lead ECG recordings of 10 seconds each from 18869 patients. Each ECG recording was annotated by cardiologists with multiple diagnostic statements, resulting in a total of 71 different ECG statements. These statements are further grouped into five high-level diagnostic categories: Norm (NORM), Myocardial Infarction (MI), ST/T Change (STTC), Conduction Disorder (CD) and Hypertrophy (HYP). In addition to the ECG waveforms, the dataset provides extensive metadata. The ECG signals are available in both high-resolution (500 Hz) and low-resolution (100 Hz) formats.

### 2.3 Training and Evaluation

The CODE dataset is used for model training following the original implementation, while the PTB XL dataset serves as an independent external dataset for evaluating both model performance and XAI behavior under domain shift. The use of PTB XL as an external dataset allows us to evaluate XAI behavior under domain shift, reflecting more realistic deployment conditions compared to in-distribution analysis on the training dataset. We replicated the 12-lead training pipeline proposed by A.H. Ribeiro *et al*. [34] using a PyTorch implementation of their 1D-ResNet architecture adapted from [19]. Specifically, we applied their preprocessing steps (resampling to 400 Hz, zero-padding each 10 s ECG to 4,096 samples) and adopted the same hyperparameter settings as in the original work. By closely following the original training protocol and changing only the deep learning framework, we ensured minimal deviations from the methods validated in [34] while demonstrating reproducibility in PyTorch, enabling the application of Captum to XAI methods [17].

In line with common practices for large-scale data mining, we used raw ECG signals without additional filtering. Our experiments revealed lower performance when filters were applied during training or prediction, corroborating evidence from prior publications that DL models can inherently handle noisy inputs [8, 14, 34]. Thus, the 1D-ResNet effectively learns diagnostic features directly from the unprocessed waveforms, leveraging its robust architecture to mitigate noise disturbances while retaining clinically valuable signal characteristics.

### 2.4 XAI Methods

We used the BenchXAI package [25] to generate relevance attributions with 15 different local post-hoc XAI methods. We applied all of these methods to the samples from the PTB XL dataset to generate relevance attributions for each sample (local) after the model was trained (post-hoc). In addition, we used the Guided Grad-CAM [38] XAI method which only works with convolutional network layers. The following XAI methods were used in this study. A shorter summary of the methods can be found in Appendix Section A.2.

#### 2.4.1 Backpropagation-based methods

Backpropagation-based XAI methods utilize the backpropagation mechanism of neural networks to propagate either the gradient of the network with respect to the input or other values, such as the network’s output. This allows them to assign attributions throughout the network, ultimately reaching the input layer.

Simonyan et al. introduced the **Input × Gradient** method in the context of image classification to generate saliency maps as explanations [44]. In this approach, they relied on the absolute values of the gradients. The use of gradients is intuitive, as gradients indicate how a change in each input dimension affects the model’s output. By multiplying the gradient by the corresponding input value, the method provides an attribution score that reflects both the input’s influence on the output and its magnitude within the input space. This method is computationally efficient, as the gradient is inherently calculated during backpropagation. However, it has limitations, as relying on gradients alone can be simplistic and may lead to misleading results by pointing to local optima of the model’s output.

Similar to Input × Gradient, the **Deconvolution** method [56] calculates the gradient of the model output with respect to the input. The primary difference between the two methods is that Deconvolution applies a rectified linear unit (ReLU) activation function to the relevance attributions during backpropagation, but only if a ReLU activation was used in the corresponding layer during the forward pass through the network.

The **Guided Backpropagation** method [45] merges the Input × Gradient [43] and Deconvolution [56] methods. It achieves this by backpropagating relevance attributions only when both the activation during the forward pass and the gradient during the backward pass are positive.

The **Guided Grad-CAM** method combines the Grad-CAM [38] and guided backpropagation methods to create more detailed and localized visual explanations. Grad-CAM generates a coarse heatmap by computing gradients of the output with respect to a specific convolutional layer. Guided backpropagation refines this heatmap by incorporating fine-grained feature information, producing higher resolution explanations that combine spatial importance with detailed gradients.

Sundararajan et al. [47] introduced **Integrated Gradients**, a method that calculates the average gradient along a straight-line path from a baseline to the input. It attributes importance to each feature by integrating these gradients, providing a more stable and theoretically grounded explanation compared to simpler gradient methods.

Bach et al. [5] introduced **Layerwise Relevance Propagation (LRP)**, a method that calculates relevance attributions by decomposing a neural network’s output layer by layer, propagating relevance backward through the network until it reaches the input. The decomposition ensures that, for any layer, the relevance attributions for all neurons sum up to the network’s output. To stabilize the method and avoid unbounded attributions caused by very small relevance attributions, they proposed adding a small stabilizing term, *ϵ* > 0, referred to as **LRP**-***ϵ***. Another stabilization method proposed in [5] is **LRP**-***α*** 1-***β*** 0, which separates positive and negative pre-activations, propagating only the positive ones. Additionally, **LRP**-***γ***, introduced in [29], assigns greater importance to positive relevance attributions.

**DeepLIFT** (Deep Learning Important FeaTures), introduced by Shrikumar et al. [42], operates similarly to the Integrated Gradients method. While Integrated Gradients calculates the average partial derivative of each feature along a straight-line path from the baseline to the input, DeepLIFT simplifies this process by approximating the same quantity in a single step, replacing the gradient at each non-linearity with its average gradient.

#### 2.4.2 Perturbation-based methods

Perturbation-based XAI methods modify an input sample to observe how these changes impact the neural network’s output. This is often done by replacing specific features or regions, such as pixels in an image, with a baseline value, typically set to zero

In the context of image classification, **Occlusion** was introduced by Zeiler et al. [56]. This method involves masking one or more features/pixels in the input, performing a forward pass through the network, and computing the difference in outputs as the relevance attribution. For image or signal data, occlusion can be applied to individual pixels or by sliding a feature mask across the image, setting all pixels within the mask to a baseline value, and calculating the resulting output difference.

**Shapley Value Sampling** was introduced by Strumbelj et al. [46] as a faster sampling based approach to calculate Shapley Values. Shapley values [39, 40] come from the field of cooperative game theory and calculate the average marginal contribution of a feature value across all possible subsets of features.

Lundberg and Lee [21] introduced SHAP (SHapley Additive exPlanations) values, which represent the Shapley values of a conditional expectation function of the original model. **GradientSHAP** [21] approximates SHAP values by computing expectations of gradients using random sampling from a distribution of baseline values, serving as an approximation of the Integrated Gradients method.

**KernelSHAP** [21], another SHAP-based method, uses the LIME [37] approach to approximate SHAP values.

Finally, **DeepLiftSHAP** [21] employs the DeepLIFT method [42] to approximate SHAP values by rewriting the DeepLIFT multipliers in terms of SHAP values (see [21] for detailed formulas).

#### 2.4.3 Surrogate methods

Surrogate XAI methods involve training interpretable models, such as linear regression, to approximate the behavior of a neural network. Due to the complexity of neural networks, these surrogate models can only approximate the network locally around a specific input sample, providing a sufficiently accurate approximation and making the surrogate model’s parameters interpretable.

Ribeiro et al. [37] introduced the **LIME** (Local Interpretable Model-agnostic Explanations) method, which trains an interpretable model locally around a specific prediction. The core concept of LIME is the fidelity-interpretability trade-off: the surrogate model must faithfully approximate the behavior of the black-box model in the local region around the input while maintaining low complexity to remain human-interpretable. As a model-agnostic approach, LIME can be applied to any AI model.

### 2.5 XAI Evaluation

#### 2.5.1 Preprocessing

Since analyzing individual ECG recordings is not only time-consuming but also does not allow for generalization, we decided to perform a quantitative analysis. First, we calculated the average heartbeats for each ECG from the PTB XL dataset by using the widely used XQRS() function of the Python WFDB package [54] for R-Peak detection and then performed a majority vote to ensure that they were detected in multiple leads filtering false detected peaks robustly. Afterward, we removed all samples with more than 10 overlapping heartbeats. The heartbeats have a length of 650ms, 250ms before the R-Peak, and 400ms after the R-Peak. For all remaining samples, the average heartbeat is then calculated as the mean overall validated heartbeats for each lead. Within each average heartbeat, we analysed four clinically relevant regions: the P-wave, QRS complex, R-peak, and T-wave. Since the actual position of these waves varies between individuals and with heart rate, clinical definitions rely on morphological rather than fixed temporal criteria [32]. For a reproducible quantitative analysis across many samples, however, fixed intervals are required. We therefore defined the following intervals based on a cardiological assessment of representative ECGs from the PTB-XL dataset, chosen to reliably contain the respective waveform in the majority of cases: P-wave from 210ms to 70ms before the R-peak (140ms), QRS complex from 60ms before to 90ms after the R-peak (150ms), R-peak region ±20ms around the R-peak, and T-wave from 180ms to 350ms after the R-peak (170ms). Figure 2 shows an example of an average heartbeat with relevant regions being highlighted.

**Figure 2:**
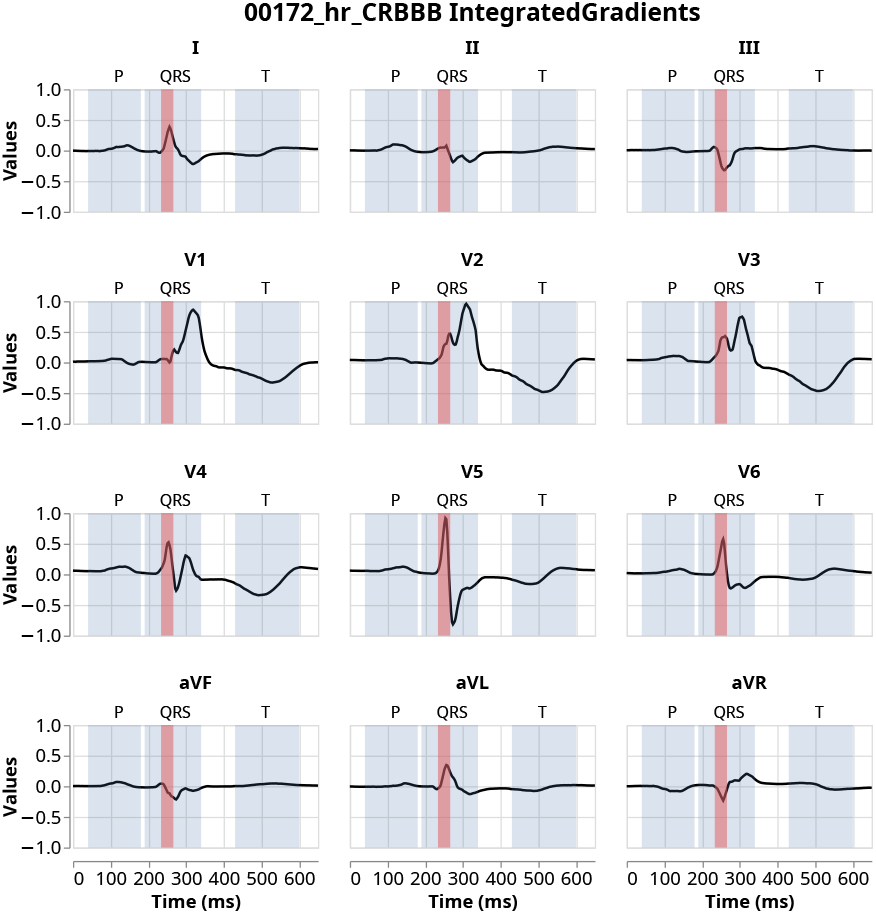
Example of average heartbeats of a patient with Complete Right Bundle Branch Block (CRBBB) with clinically relevant P-wave, QRS complex, R-peak region (red), and T-wave regions highlighted.

The corresponding relevances for all heartbeats are then identically averaged to create average relevance attributions for each average heartbeat.

Since the ranges of relevance attributions vary greatly across different post-hoc XAI methods, we decided to normalize each average ECG relevance attribution using the same rank normalization employed in [25]. This will normalize all relevance attributions into the range [−1, 1] while keeping the correct order of relevance attributions within each averaged ECG.

After normalizing the relevance attributions, we decided to focus our further evaluation on lead V1 instead of on all twelve leads. From a clinical standpoint, Lead V1 is often emphasized because of its anatomic orientation typically captures a strong signal from the right atrium and right ventricle. This proximity makes V1 especially sensitive to conduction and rate abnormalities, enabling clearer detection of the characteristic waveforms and timing changes pertinent to the six target arrhythmias [48].

Focusing on a single lead further enables a more concise and interpretable analysis of the XAI methods. However, we acknowledge that this choice may limit the generalizability of our observations across all leads, and extending the analysis to additional leads represents an important direction for future work.

To evaluate the normalized XAI relevance attributions for the four clinically relevant regions, we first plotted them using box-plots to gauge their overall distribution for each XAI method and for all correctly classified samples for each disease.

#### 2.5.2 Statistical Analysis

Afterward, we performed a one-sided Wilcoxon signed-rank test [41, 52] to analyse whether the mean relevance of a clinically relevant region was significantly larger than that of each of the other regions. Let *X* be the mean relevance of one region of the ECG (e.g., P-Wave) and let *Y* be the mean relevance of another region of the same ECG (e.g., QRS-Complex). The two-sided null hypothesis can be written as *H*_0_ : „median (*X*) = median (*Y*) “meaning there is no difference between the medians of the two distributions with *H*_1_ : „median (*X*) ≠ median (*Y*)”. The one-sided test tests whether one distribution is significantly larger than the other. We then plotted the p-values of these tests in a heatmap to show which region had significantly larger attributions than another region. A significant p-value (<0.05) indicates the rejection of the null hypotheses for a specific XAI method. Because of repeated testing for each clinically relevant region, XAI method, and disease, all p-values were corrected using the Bonferroni correction [11] to reduce the likelihood of making a Type-1-error (incorrectly rejecting *H*_0_).

Lastly, we performed a one-sided Mann–Whitney U test [41, 52] to test for significant differences in relevance attributions of the clinically relevant regions between correctly and incorrectly classified ECGs, more specifically between true positives and false positives. Let *X* and *Y* be the relevances of two different ECGs but for the same region (e.g., QRS-Complex). A general formulation of the two-sided null hypothesis can be expressed as *H*_0_ : „median (*X*) = median (*Y*) “meaning the median of the relevances *X* represents equals the median of the relevances *Y* represents with an alternative hypothesis *H*_1_ : „median (*X*) ≠ median (*Y*)”. The one-sided version of this test tests whether one sample distribution is significantly larger or smaller than the other. A significant p-value (<0.05) indicates the rejection of the null hypotheses for a specific region and XAI method. Because of repeated testing for each clinically relevant region, XAI method, and disease, all p-values were corrected using the Bonferroni correction [11].

## 3 Results

First, we analyzed the performance of our trained model on the PTB XL dataset. As our XAI analysis focuses on correctly identified disease cases, sensitivity is the primary performance metric of interest, complemented by TP and FP counts to provide additional context on model behavior. Overall, our trained model had high sensitivity scores of > 0.9 for most diseases, except 1AVB and SBRAD, while AFIB and SBRAD had the fewest false positives (see Table 1).

**Table 1:** Table with information on PTB-XL dataset after preprocessing according to Section 2.5.1. It summarizes descriptive performance metrics across disease classes.

| Disease | Total Samples | TP | FP | Sensitivity |
| --- | --- | --- | --- | --- |
| 1AVB | 562 | 442 | 220 | 0.79 |
| AFIB | 994 | 807 | 85 | 0.91 |
| CRBBB | 375 | 363 | 257 | 0.99 |
| CLBBB | 292 | 278 | 344 | 0.99 |
| SBRAD | 506 | 315 | 28 | 0.62 |
| STACH | 685 | 636 | 132 | 0.96 |

When looking at the overall distributions of normalized relevance attributions for CRBBB (see Figure 3) we found that the LIME method produced mainly zero attributions with only a few outliers, while the LRP methods (LRP, LRP-*ϵ*, LRP-*γ*, and LRP-*α*1-*β*0) had median attributions of zero, with a larger interquartile range. Moreover, KernelShap, LIME, and the LRP methods showed almost no difference in median attributions between all four clinically relevant regions (P-wave, QRS complex, R-Peak region, T-wave). We found that this was also true for the other five diseases (see Appendix Section A.3). Since these six XAI methods were not able to sufficiently discern between the relevant regions for all diseases, we have decided to disregard them for the remainder of the analysis.

**Figure 3:**
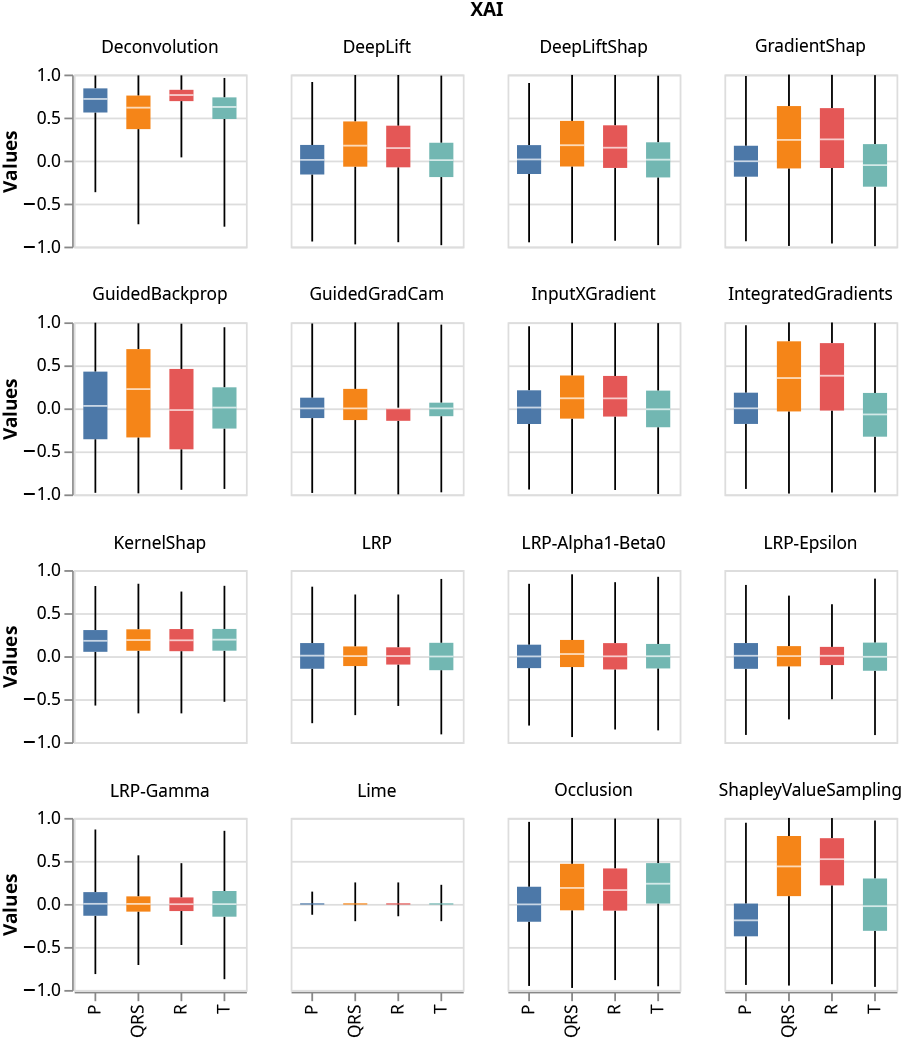
Boxplots showing the distribution of normalised relevance attributions for all true positive CRBBB samples for all 16 post-hoc XAI methods and all four clinically relevant regions.

To validate how well or even if our model learned clinically relevant features for each disease we used a one-sided wilcoxon signed-rank test to observe whether the clinically relevant features had significantly higher relevance attributions then the other three features (p-value < 0.05, see Section 2.5.2). We then performed a majority vote over the results for all remaining ten XAI methods to get a more robust decision on whether the model learned the correct features for each disease. Heatmaps with p-values for each disease, feature, and XAI method can be found in Appendix Section A.4. For 1AVB, AFIB the P-wave is the most relevant feature, while for CRBBB and CLBBB, the QRS complex is the most relevant feature. We did not consider the diseases SBRAD and STACH for this analysis, since they are rhythm based diagnoses, which are typically diagnosed across multiple heartbeats (and we are currently looking at an averaged heartbeat). Conduction and rate abnormalities – such as 1AVB, AFIB – primarily originating in the atria affect the morphology or timing of the P-wave, while conduction blocks located in the ventricles like CRBBB and CLBBB predominantly alter the QRS complex. These ECG features align with the standard interpretation guidelines [48] as key diagnostic signatures for each respective arrhythmia.

In Table 2, we can see that based on the majority vote of all ten XAI methods, our model was able to correctly learn clinically relevant features for AFIB, CRBBB. For these diseases, the relevant features had significantly larger relevance attributions more than 50% of the time. Additionally, the results from Table 2 clearly show that the XAI methods DeepLift, DeepLiftShap, and Occlusion were the only ones able to correctly identify the clinically relevant features over all four diseases, while Guided Backpropagation and Integrated Gradients only failed to do so for CLBBB and AFIB respectively.

**Table 2:** Table showing counts on how often each relevant feature had significantly larger relevance attributions than the other features (p-value < 0.05) for 1AVB, AFIB, CRBBB, and CLBBB and all XAI methods. Since there are three other features in total and ten XAI methods, a total of 30 significantly larger p-values could have been achieved for each disease. The percentage value used for the majority vote is calculated by dividing the total count for each disease by 30. A positive majority vote (>50%) is highlighted.

| XAI | 1AVB | AFIB | CLBBB | CRBBB |
| --- | --- | --- | --- | --- |
| Deconvolution | 3 | 2 | 0 | 0 |
| DeepLift | 1 | 2 | 2 | 2 |
| DeepLiftShap | 1 | 2 | 3 | 3 |
| GradientShap | 0 | 0 | 0 | 2 |
| GuidedBackprop | 1 | 3 | 0 | 3 |
| GuidedGradCam | 0 | 2 | 0 | 2 |
| InputXGradient | 0 | 2 | 0 | 2 |
| IntegratedGrad. | 1 | 0 | 3 | 2 |
| Occlusion | 2 | 3 | 2 | 1 |
| ShapleyValueS. | 0 | 0 | 3 | 2 |
| Total | 9 | 16 | 13 | 19 |
| Percent | 30% | 53.33% | 43.33% | 63.33% |

Lastly, we wanted to analyze whether there are significant differences in relevance attributions between true positive and false positive classified samples for each disease. To test this, we performed a one-sided Mann–Whitney U test (see Section 2.5.2).

When looking at the heatmaps in Figure 4, we would have expected significant differences (p-value < 0.05) in P-wave attributions for 1AVB, AFIB. However, there was only a majority (6 out of 10 XAI methods) of significant differences for AFIB. For SBRAD and STACH most significant differences were found in the P-Wave region, even though we did not expect to find any significant differences for these diseases since they are clinically diagnosed over multiple hearbeats and not an average heartbeat, which the XAI methods are looking at. However, since the underlying AI model is trained on the full length ECG, it could indicate that the model learned the distances between P-Waves of multiple heartbeats. For SBRAD, it should be noted that there were only 28 false positive samples (see Table 1), which could explain the lack of significant p-values. For CLBBB and CRBBB, there should be a majority of significant differences in the QRS complex, but in our case, this is only true for CRBBB (8 out of 10 XAI methods). The underlying ECG signals (see Figure 4 top) show representative examples for all six diseases. In 1AVB, the attributions highlight the P-wave where the electric impulse is delayed. For AFIB, the absence of the P-wave is indicated by the attributions. In the bundle branch blocks (CLBBB and CRBBB), the broad QRS is visible and shows high attributions. For the rhythm disorders (SBRAD, STACH) the attributions are evenly spaced across the averaged heartbeat, because these disorders are typically not diagnosed over an averaged heartbeat, but over multiple heartbeats.

**Figure 4:**
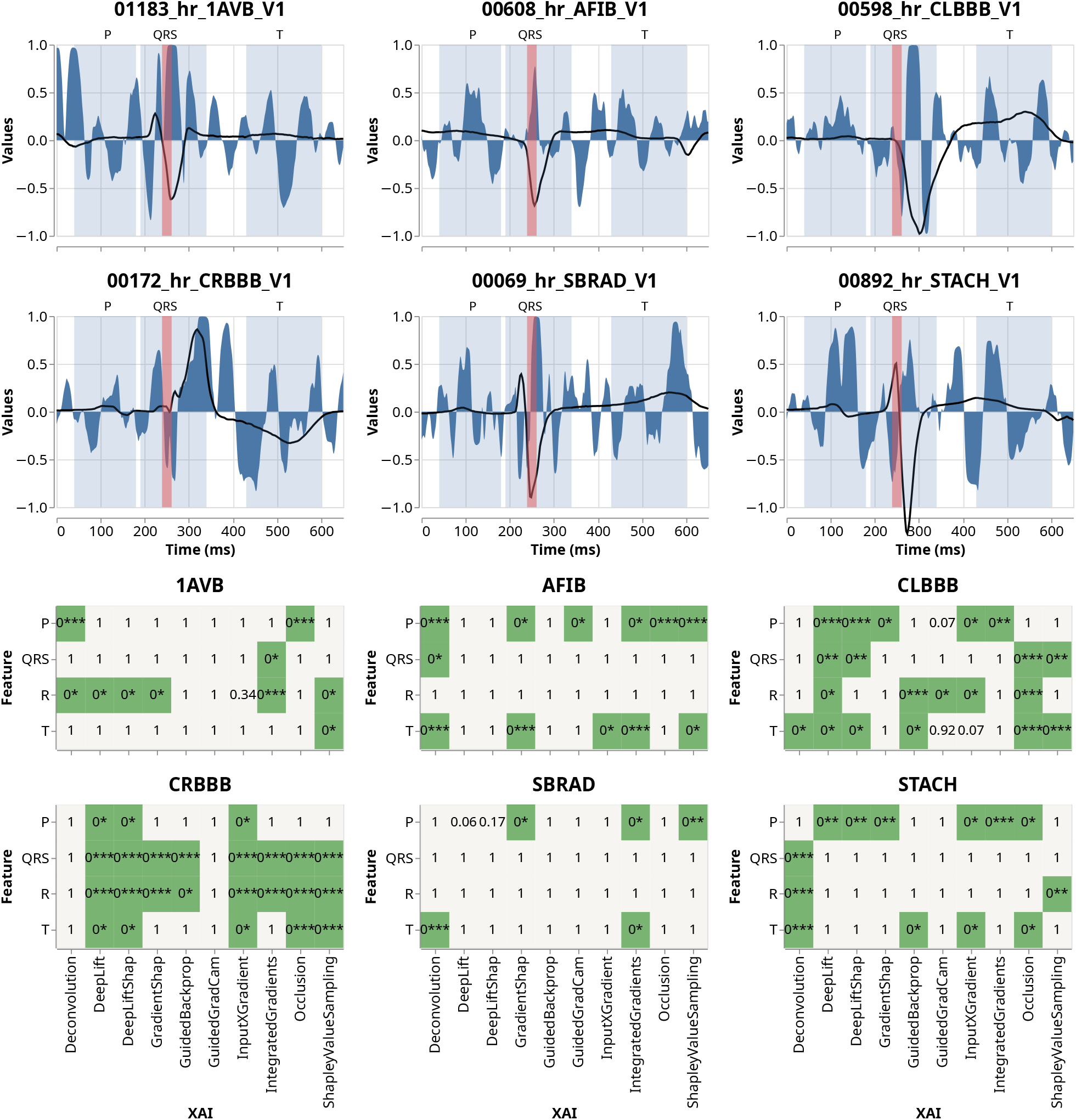
Top: Exemplary averaged heartbeats from lead V1 for each disease class. The P-wave, QRS complex, R-peak region (red), and T-wave are indicated as physiologically relevant segments. Integrated Gradients attributions are overlaid as area plots to visualize feature relevance across the cardiac cycle. Bottom: Heatmaps of statistical significance (one-sided Mann–Whitney U test) comparing relevance attributions between true positive and false positive samples across features, XAI methods, and disease classes. Significant results indicate rejection of the null hypothesis that both distributions are identical in favor of higher relevance in true positives. P-values are Bonferroni-corrected for multiple comparisons. To improve readability across very small values, p-values are grouped into magnitude bins and displayed using the notation 0*, 0**, and 0*** corresponding to p < 0.05, p < 1e-20, and p < 1e-50, respectively.

## 4 Discussion

In this study, we demonstrated the potential of gaining more robust explanations on the decision-making process of a DL model by using an ensemble approach of post-hoc XAI methods. Additionally, we showed the importance of selecting appropriate XAI methods for a specific classification task, as well as their capabilities to identify false positive samples. Rather than providing a clinical decision-support tool, our work is intended to support the informed selection and interpretation of XAI methods when applied to ECG classification models.

We found that LRP-type XAI methods require careful parameter tuning to produce stable and interpretable relevance attributions, as well as KernelShap and LIME. A possible explanation for this may lie in the structural differences between image time-series data. While these methods have been widely validated in computer vision tasks, ECG signals have strong temporal dependencies that do not match well with the patch- or pixel-based approaches used by many of these methods. In particular, perturbation-based methods such as LIME and KernelSHAP may generate out-of-distribution samples when applied to temporal data, potentially leading to unstable or physiologically implausible attributions [9, 18, 23, 51]. Improvements of these methods in this domain is an active field of research, with many adaptations being made to the algorithm [2, 23, 24]. For the LRP methods, this behavior is also noticed by other researchers [16, 25]. As suggested by Kohlbrenner et al. [16], using different LRP methods for different neural network layers in combination could improve their performance.

Our ensemble majority vote approach using the XAI methods Deconvolution, DeepLift, DeepLiftShap, GradientShap, Guided Backpropagation, GuidedGradCam, Input×Gradient, Integrated Gradients, Occlusion, and ShapleyValueSampling, suggest that the model captures features that are consistent with clinically relevant patterns for AFIB, CRBBB. This majority vote approach gives us more robust explanations since post-hoc XAI methods by themselves have been shown to be very sensitive to small changes in the data or model [20, 55]. When looking at the XAI methods by themselves, only DeepLift, DeepLiftShap, and Occlusion showed that our model might have learned clinically relevant features for all diseases.

Additionally, our ensemble approach showed that for our model and dataset, the 10 remaining XAI methods could potentially be employed to discern between true positive and false positive classified samples for diseases AFIB and CRBBB. This capability highlights the potential of XAI methods as diagnostic tools for model inspection and error analysis, particularly in distinguishing false positive from true positive predictions. These differences can not only improve our understanding of the model’s strengths and weaknesses but also support the development of more transparent, reliable, and robust AI systems.

We note that the number of false positive samples in the SBRAD class is limited (n = 28), which may reduce the statistical power and increase uncertainty in subgroup-specific statistical comparisons. Additionally, the use of fixed temporal intervals represents a limitation of our approach, and future work will explore morphology-based segmentation to better capture inter-individual variability. Especially for rhythm diagnoses such as SBRAD and STACH, which are defined by R–R interval across multiple heartbeats, restricting the analysis to an averaged heartbeat may limit the ability of XAI methods to identify meaningful relevance.

The comparison to other studies demonstrates that the performance of XAI methods might differ between various DL models since methods like GradCam, which underperforms in our experimental setting, are successfully employed with multiple datasets and DL models in other ECG classification tasks [3, 33]. While our comprehensive evaluation is based on one of the largest ECG datasets, multiple diseases, and a variety of XAI methods, the lack of additional datasets and DL architectures demonstrates a limitation of our current study. Therefore, evaluating the proposed ensemble post-hoc XAI framework in more external datasets and the integration of additional DL architectures is an avenue for future work.

Lastly, we acknowledge that the several gradient-based XAI methods in our ensemble may produce correlated attributions. However, the ensemble majority-vote approach is intended as a robustness strategy rather than a statistically independent aggregation, and exploring inter-method similarity remains an important direction for future work. Future work could also explore alternative ensemble aggregation strategies beyond the majority-vote threshold (≥50%), which was used in this study as a simple and interpretable criterion ensuring that only features supported by more than half of the XAI methods are considered robust.

In summary, we show the importance of using ensemble XAI approaches to enhance the interpretability and reliability of DL models, supporting the development of more transparent and interpretable AI systems for healthcare applications. Our results indicate that no single XAI method consistently outperforms others across all settings; instead, ensemble-based interpretation or careful task-dependent selection may provide more reliable insights.

## 5 Conclusion

This study demonstrates the effectiveness of ensemble post-hoc XAI methods in improving the interpretability, robustness, and clinical relevance of deep learning models for ECG classification. By aggregating outputs from multiple explanation techniques, we were able to identify consistent, meaningful patterns associated with different heart abnormalities.

The results emphasize the importance of method selection and parameter tuning in XAI applications, and show that ensemble-based strategies can mitigate the limitations of individual explanation methods. In particular, the ability of our ensemble approach to detect false positives offers promising opportunities for model debugging and refinement.

Future research should focus on applying this ensemble XAI framework to different datasets and model architectures to evaluate its generalizability. Expanding the diversity of test conditions will help determine the reliability and practical applicability of ensemble explanations in real-world clinical settings.

Ultimately, our findings support the integration of ensemble XAI techniques as a key component in the development of trustworthy and transparent AI systems for healthcare.

## 6 Declarations

### 6.1 Ethics approval and consent to participate

Not applicable.

### 6.2 Consent for publication

Not applicable.

### 6.3 Availability of data and materials

The data that support the findings of this study are available from https://doi.org/10.17044/scilifelab.15169716.v1 but restrictions apply to the availability of these data, which were used under license for the current study, and so are not publicly available. Data are however available from the authors upon reasonable request and with permission of https://doi.org/10.17044/scilifelab.15169716.v1. The code for our analysis is available under https://github.com/HauschildLab/EnsembleXAIForTime.

### 6.6 Authors’ contributions

A-C.H., N.S., P.H. and J.M.M conceptualised the project. M.C.M. was responsible for the data acquisition. P.H. implemented the training of the AI model. J.M.M. performed all the XAI analysis. J.M.M. and P.H. generated the figures and prepared the results. P.H. and K.E.S. validated the results clinically. J.M.M., P.H., M.C.M., and K.E.S. wrote the manuscript. All authors performed review and editing.

## 6.7 Acknowledgements

This work used the Scientific Compute Cluster at GWDG, the joint data centre of Max Planck Society for the Advancement of Science (MPG) and University of Göttingen.

## A Appendix

### A.1 Data

**Table 3:** Comparison of cohort statistics and ECG abnormalities between CODE and PTB-XL. Values are *n* (%) unless otherwise specified.

|  | CODE | PTB-XL (All) |
| --- | --- | --- |
| **Number of records** | 2,322,513 | 21,837 |
| **Abnormality** |  |  |
| 1dAVb | 35,759 (1.5%) | 562 (2.6%) |
| CRBBB | 63,528 (2.7%) | 375 (1.7%) |
| CLBBB | 39,842 (1.7%) | 292 (1.3%) |
| SBRAD | 37,949 (1.6%) | 506 (2.3%) |
| AFIB | 41,862 (1.8%) | 994 (4.6%) |
| STACH | 49,872 (2.1%) | 685 (3.1%) |
| **Sex** |  |  |
| Male | 922,780 (39.7%) | 11,162 (51.1%) |
| Female | 1,399,733 (60.3%) | 10,675 (48.9%) |
| **Age (years)** |  |  |
| Mean | 51.2 | 62.3 |

### A.2 XAI Methods

**Table 4:**
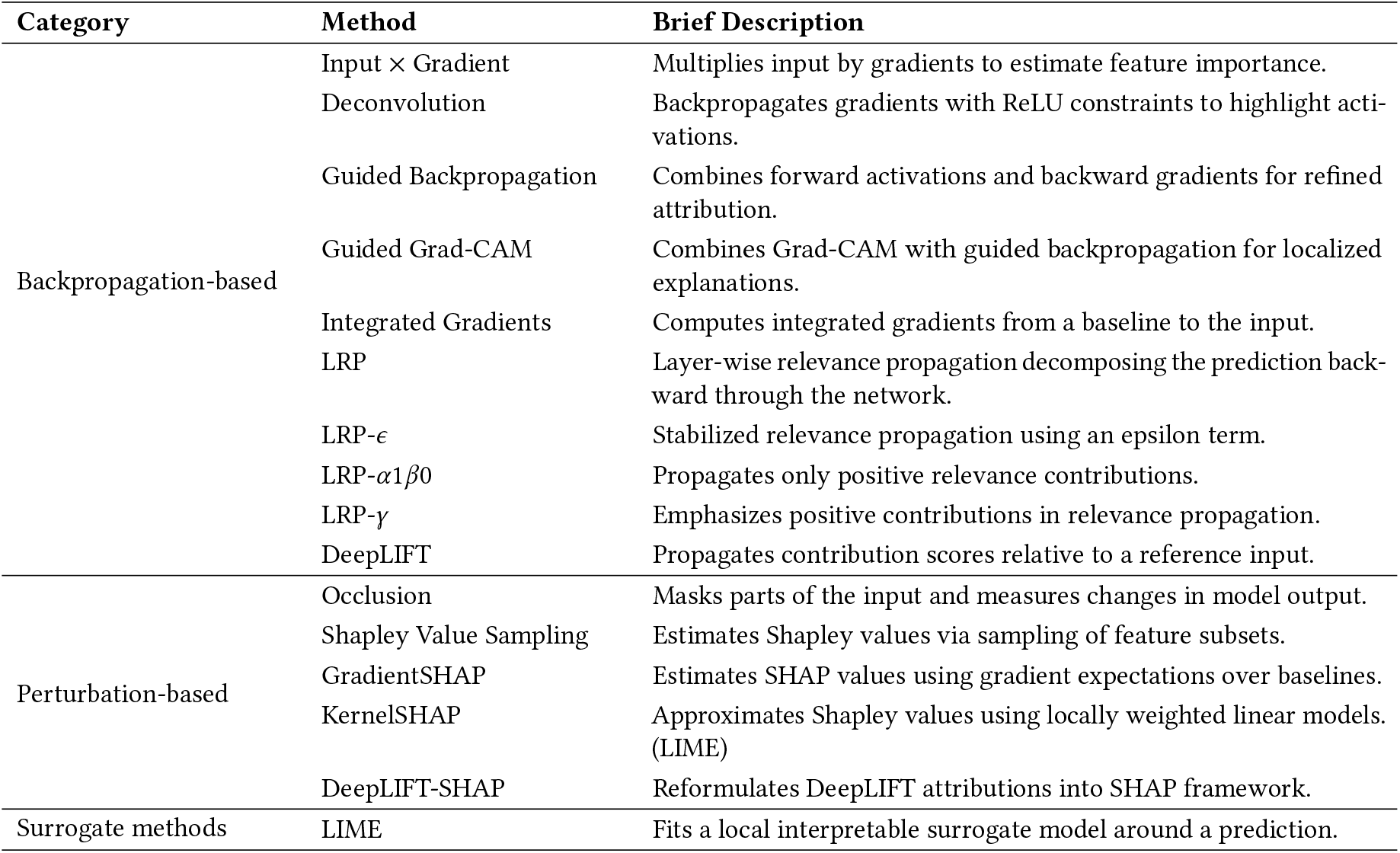
Overview of the 16 XAI methods used in this study, grouped by methodological category.

### A.3 Boxplots

**Figure 5:**
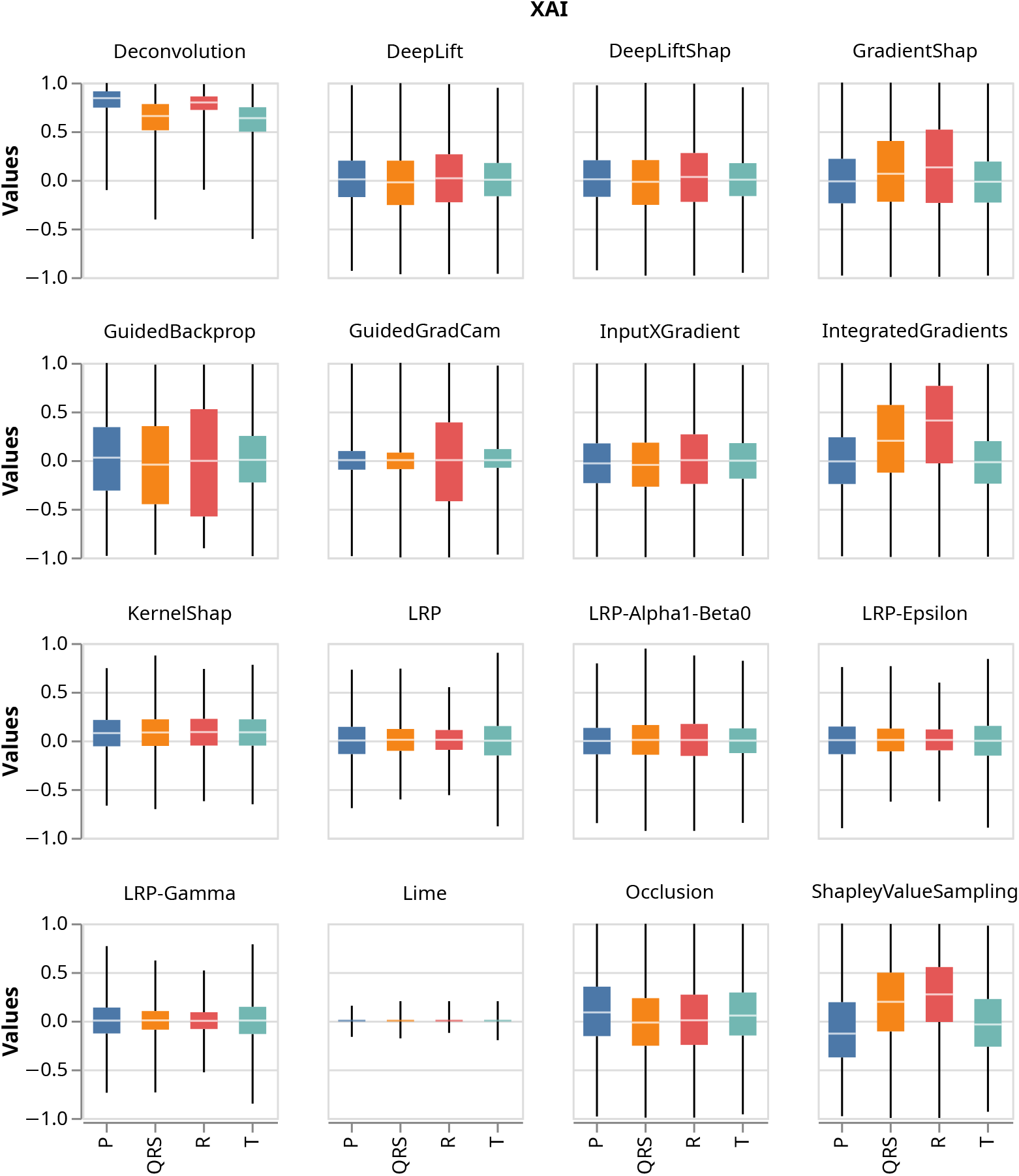
Boxplots showing distribution of normalised relevance attributions for all true positive 1AVB samples for all 16 post-hoc XAI methods and all four clinically relevant regions.

**Figure 6:**
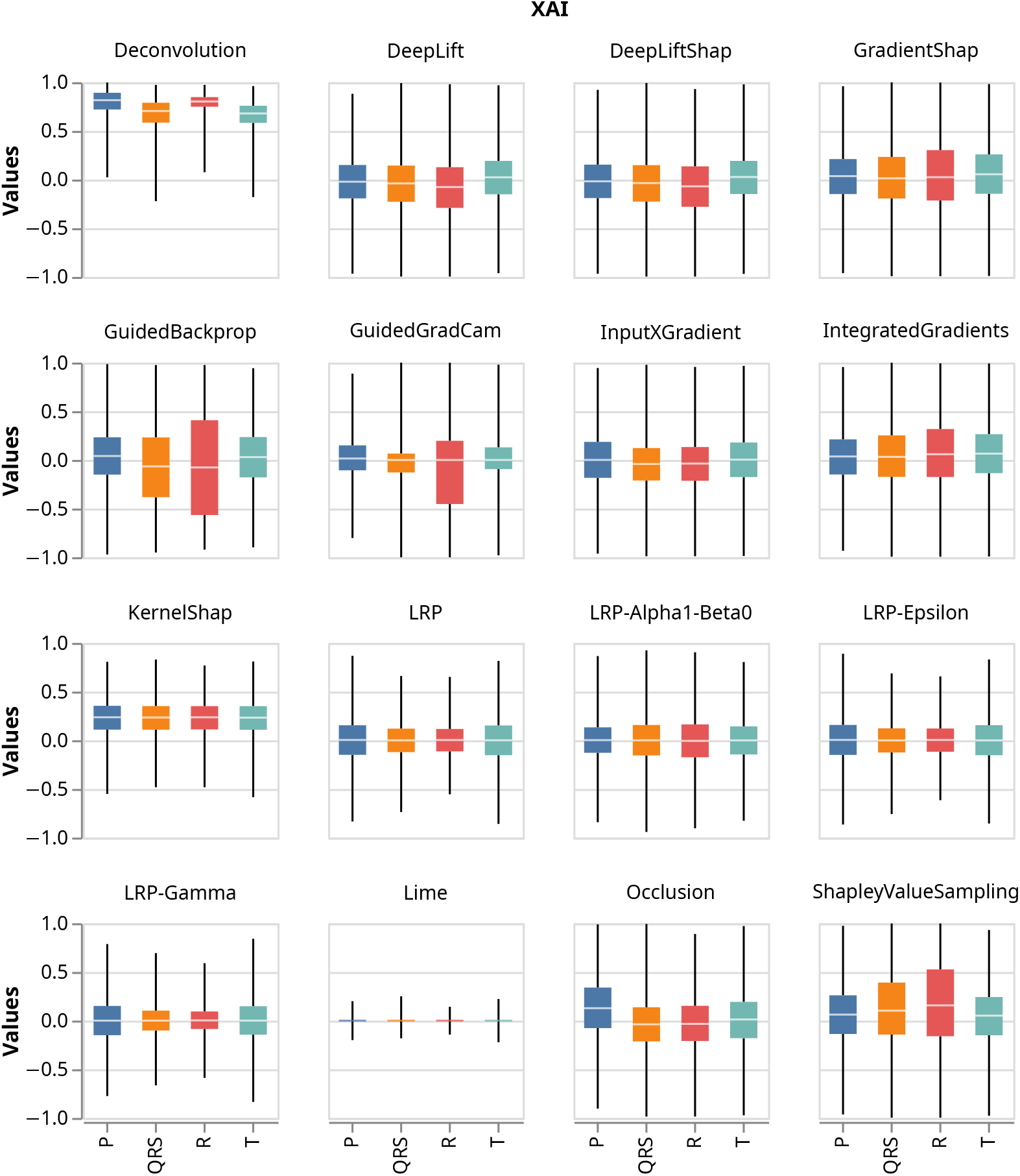
Boxplots showing distribution of normalised relevance attributions for all true positive AFIB samples for all 16 post-hoc XAI methods and all four clinically relevant regions.

**Figure 7:**
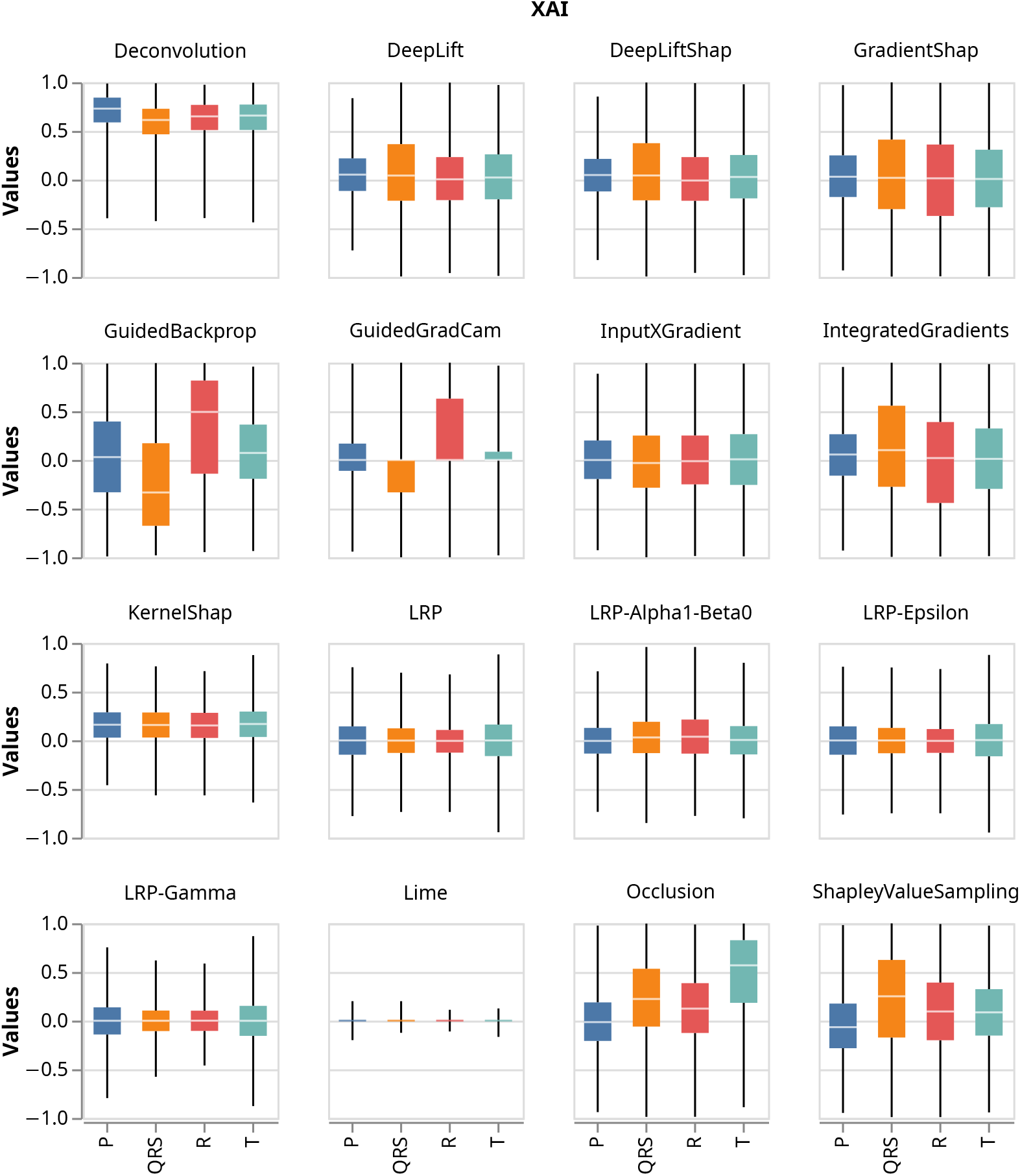
Boxplots showing distribution of normalised relevance attributions for all true positive LBBB samples for all 16 post-hoc XAI methods and all four clinically relevant regions.

**Figure 8:**
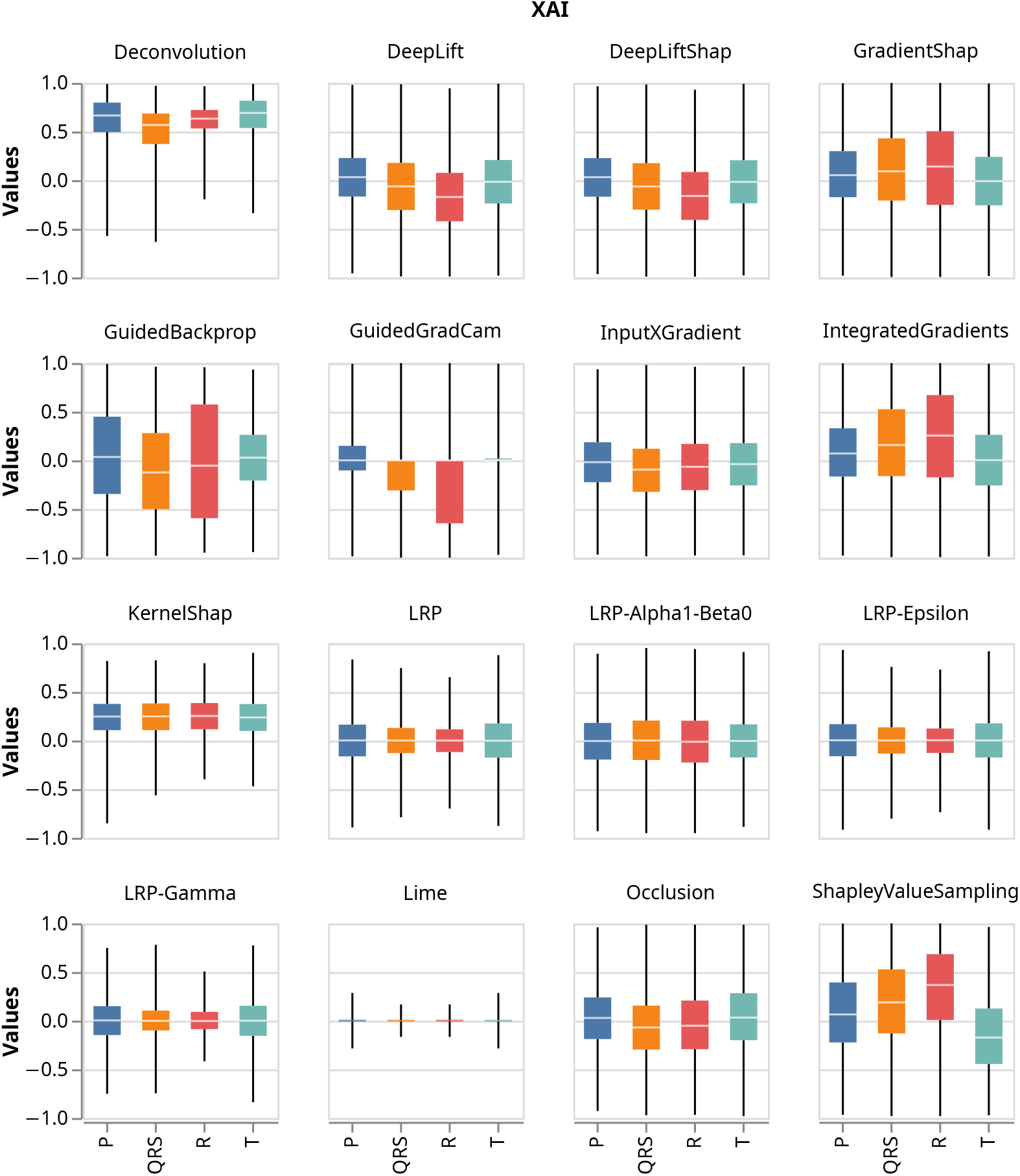
Boxplots showing distribution of normalised relevance attributions for all true positive SBRAD samples for all 16 post-hoc XAI methods and all four clinically relevant regions.

**Figure 9:**
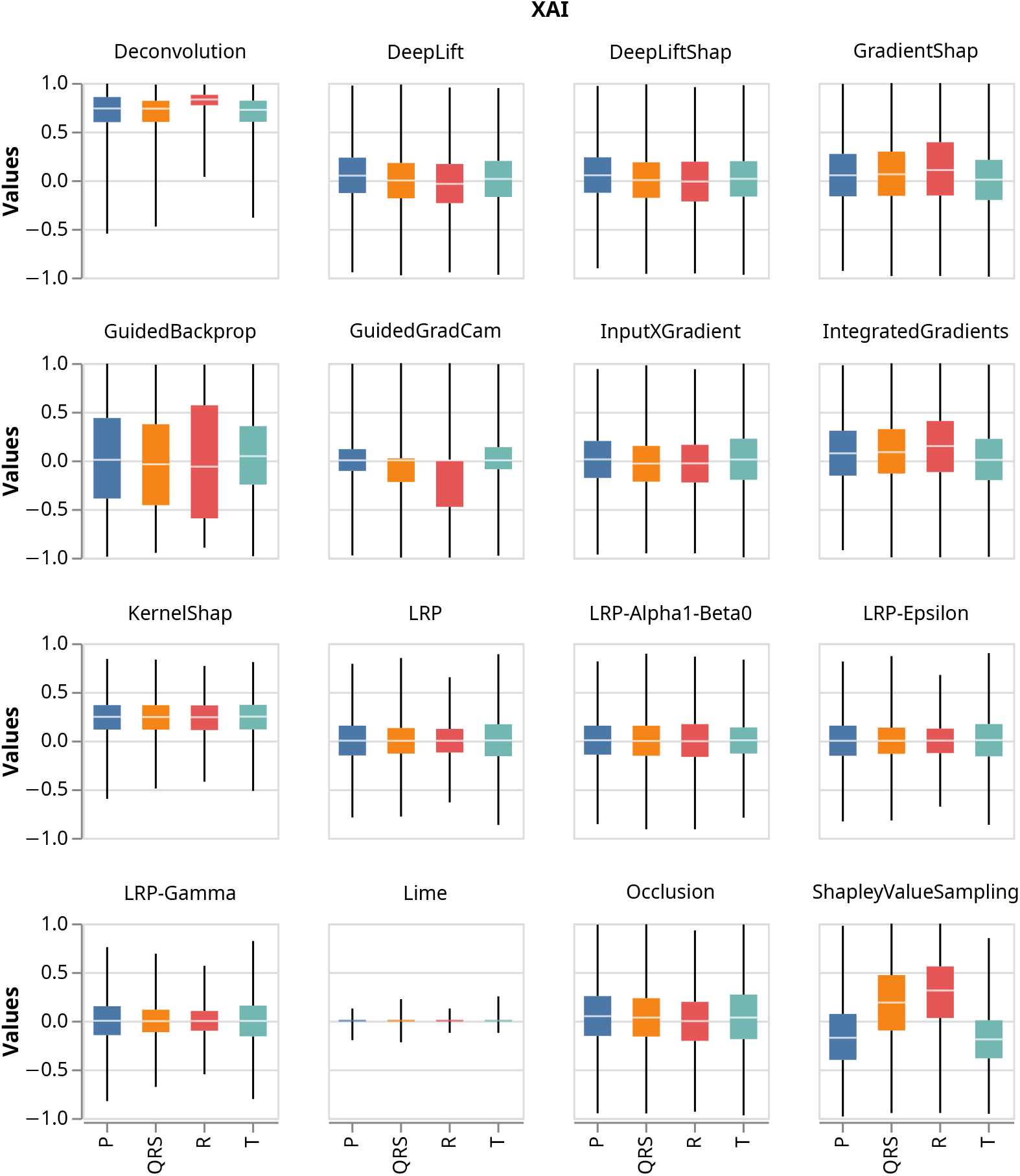
Boxplots showing distribution of normalised relevance attributions for all true positive STACH samples for all 16 post-hoc XAI methods and all four clinically relevant regions.

### A.4 Wilcoxon

**Figure 10:**
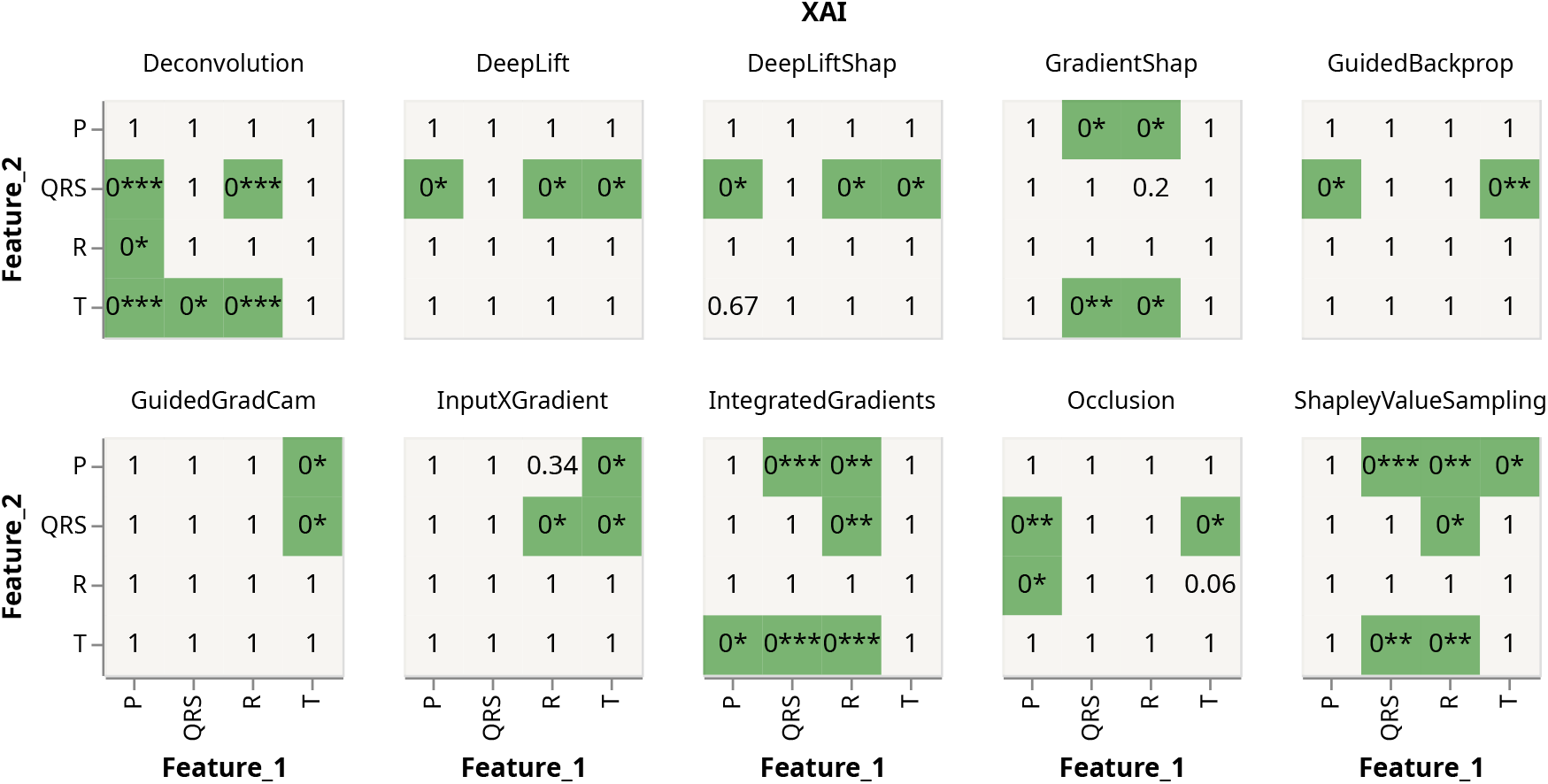
Heatmaps showing significant p-values (<0.05) when performing a one sided Wilcoxon test between all features for all ten XAI methods on all true positive 1AVB samples. A significant p-value indicates the rejection of *H*_0_ : median(mean(Relevance region 1) - mean(Relevance region 2)) = 0 in favor of *H*_1_ : median(mean(Relevance region 1) - mean(Relevance region 2)) > 0, indicating that feature 1 is significantly larger than feature 2. P-values were corrected using Bonferroni correction to account for multiple testing. 0***: p-value < 1e-50, 0**: p-value < 1e-20, 0*: p-value < 0.05.

**Figure 11:**
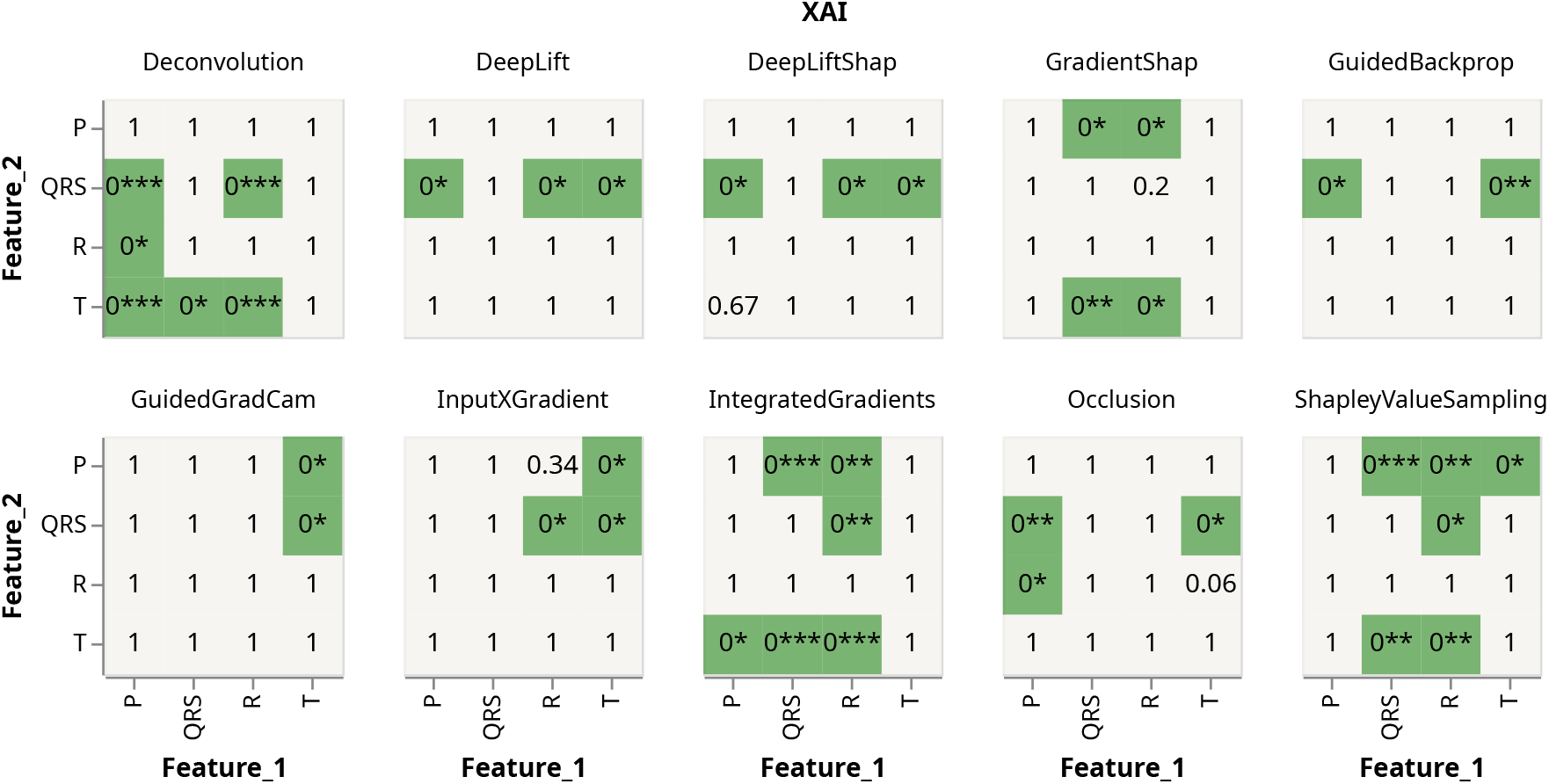
Heatmaps showing significant p-values (<0.05) when performing a one sided Wilcoxon test between all features for all ten XAI methods on all true positive AFIB samples. A significant p-value indicates the rejection of *H*_0_ : median(mean(Relevance region 1) - mean(Relevance region 2)) = 0 in favor of *H*_1_ : median(mean(Relevance region 1) - mean(Relevance region 2)) > 0, indicating that feature 1 is significantly larger than feature 2. P-values were corrected using Bonferroni correction to account for multiple testing. 0***: p-value < 1e-50, 0**: p-value < 1e-20, 0*: p-value < 0.05.

**Figure 12:**
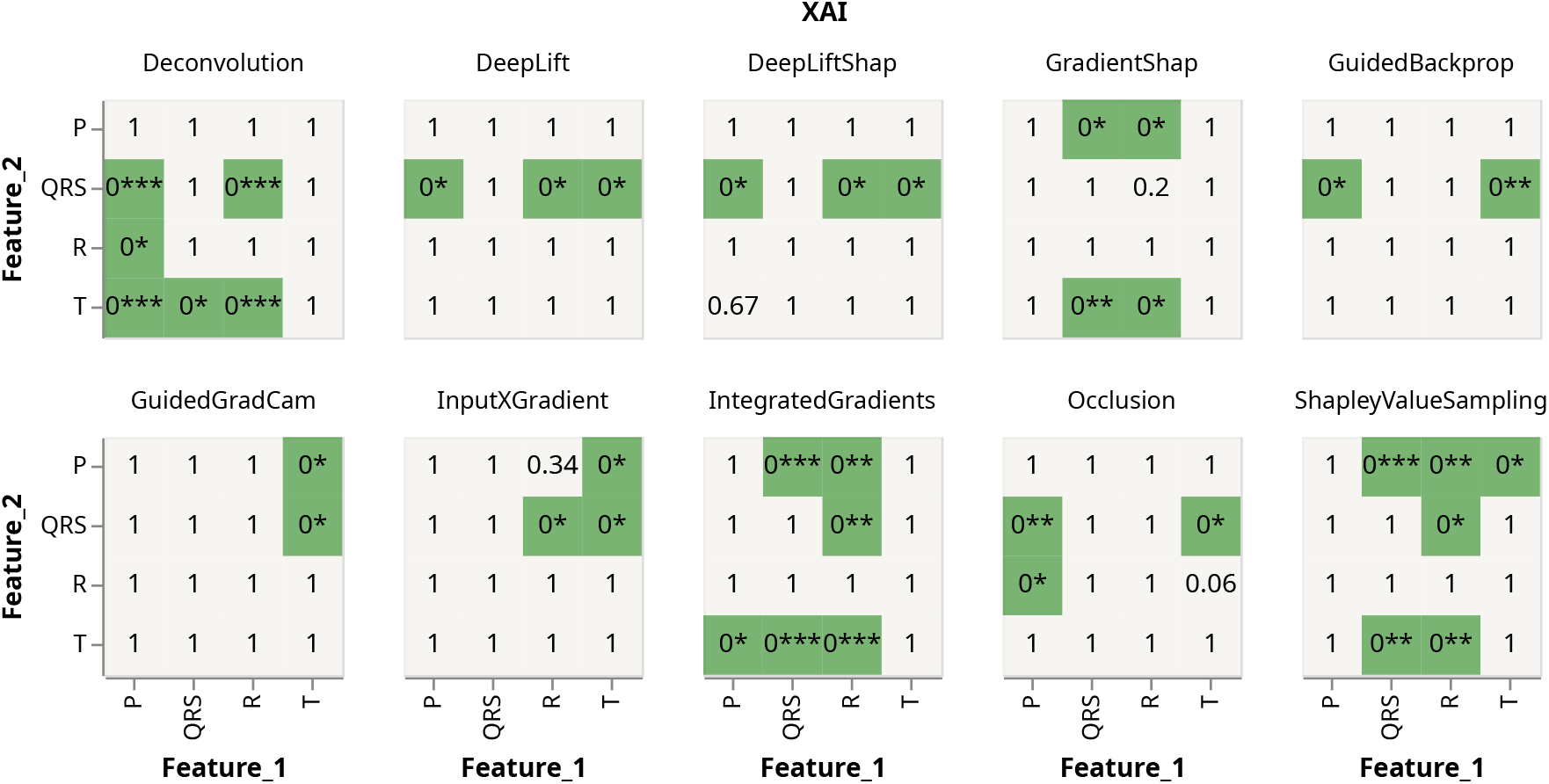
Heatmaps showing significant p-values (<0.05) when performing a one sided Wilcoxon test between all features for all ten XAI methods on all true positive LBBB samples. A significant p-value indicates the rejection of *H*_0_ : median(mean(Relevance region 1) - mean(Relevance region 2)) = 0 in favor of *H*_1_ : median(mean(Relevance region 1) - mean(Relevance region 2)) > 0, indicating that feature 1 is significantly larger than feature 2. P-values were corrected using Bonferroni correction to account for multiple testing. 0***: p-value < 1e-50, 0**: p-value < 1e-20, 0*: p-value < 0.05.

**Figure 13:**
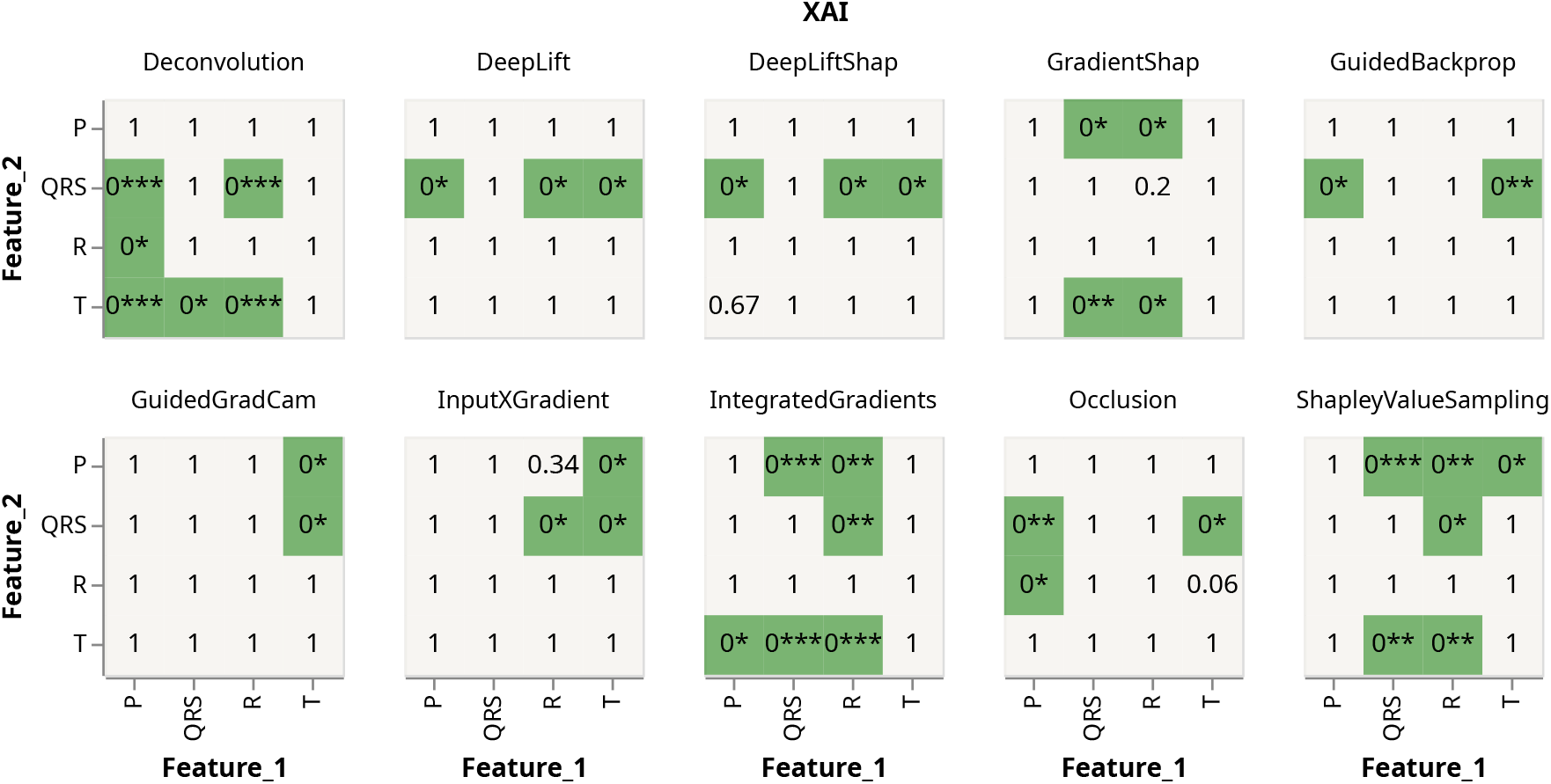
Heatmaps showing significant p-values (<0.05) when performing a one sided Wilcoxon test between all features for all ten XAI methods on all true positive RBBB samples. A significant p-value indicates the rejection of *H*_0_ : median(mean(Relevance region 1) - mean(Relevance region 2)) = 0 in favor of *H*_1_ : median(mean(Relevance region 1) - mean(Relevance region 2)) > 0, indicating that feature 1 is significantly larger than feature 2. P-values were corrected using Bonferroni correction to account for multiple testing. 0***: p-value < 1e-50, 0**: p-value < 1e-20, 0*: p-value < 0.05.

